# Genome-wide transcriptome analysis identifies alternative splicing regulatory network and key splicing factors in mouse and human psoriasis

**DOI:** 10.1101/259291

**Authors:** Jin Li, Peng Yu

**Affiliations:** Department of Electrical and Computer Engineering, College Station, TX 77843, USA; TEES-AgriLife Center for Bioinformatics and Genomic Systems Engineering, Texas A&M University, College Station, TX 77843, USA

**Keywords:** Alternative splicing, Splicing factor, Regulatory network, Psoriasis

## Abstract

Psoriasis is a chronic inflammatory disease that affects the skin, nails, and joints. For understanding the mechanism of psoriasis, though, alternative splicing analysis has received relatively little attention in the field. Here, we developed and applied several computational analysis methods to study psoriasis. Using psoriasis mouse and human datasets, our differential alternative splicing analyses detected hundreds of differential alternative splicing changes. Our analysis of conservation revealed many exon-skipping events conserved between mice and humans. In addition, our splicing signature comparison analysis using the psoriasis datasets and our curated splicing factor perturbation RNA-Seq database, SFMetaDB, identified nine candidate splicing factors that may be important in regulating splicing in the psoriasis mouse model dataset. Three of the nine splicing factors were confirmed upon analyzing the human data. Our computational methods have generated predictions for the potential role of splicing in psoriasis. Future experiments on the novel candidates predicted by our computational analysis are expected to provide a better understanding of the molecular mechanism of psoriasis and to pave the way for new therapeutic treatments.

## Introduction

Psoriasis is a chronic inflammatory skin disease with symptoms of well-defined, raised, scaly, red lesions on skin. It is characterized by excessive growth and aberrant differentiation of epidermal keratinocytes. A number of known psoriasis susceptibility loci have been identified^1^, some of which are shared with other chronic inflammatory diseases^2^. Psoriasis also shares pathways with other diseases. For example, the interleukin-23 (IL-23) pathway and nuclear factor-κB (NFκB) pathway are associated with psoriasis^3^, while the IL-23 pathway is a therapeutic target of Crohn’s disease^4^, and the dysregulation in the NFκB pathway contributes to Huntington’s disease^5^. Despite great progress made over the past few years, the exact causes of psoriasis remain unknown^6^.

To discover the disease mechanisms, significant effort has been devoted to analyzing psoriasis gene expression. For example, in a study of small and large plaque psoriasis, microarray gene expression analysis revealed the up-regulation of genes in the IL-17 pathway in psoriasis. But the expression of genes in this pathway of small plaque psoriasis is significantly higher than that of large plaque psoriasis, and negative immune regulators like CD69 and FAS have been found to be down-regulated in large plaque psoriasis. This result suggests that the down-regulation of these negative immune regulators contributes to the molecular mechanism of large plaque psoriasis subtypes^7^.

As high-throughput sequencing becomes the mainstream technology, RNA-Seq has also been used for measuring gene expression to gain biological insights of psoriasis. For example, a recent RNA-Seq‒based gene expression study of a large number of samples from lesional psoriatic and normal skin uncovered many differentially expressed genes in immune system processes^8^. The co-expression analysis based on this dataset detected multiple co-expressed gene modules, including a module of epidermal differentiation genes and a module of genes induced by IL-17 in keratinocytes. This study also discovered key transcription factors in psoriasis and highlighted the processes of keratinocyte differentiation, lipid biosynthesis, and the inflammatory interaction among myeloid cells, T-cells, and keratinocytes in psoriasis.

The high resolution of RNA-Seq data allows for study of not only gene expression but also splicing in psoriasis. A recent analysis of psoriasis RNA-Seq data revealed around 9,000 RNA alternative splicing isoforms as a significant feature of this disease^9^. Another study showed that serine/arginine-rich splicing factor 1 (SRSF1) promoted the expression of type-I interferons (IFNs) in psoriatic lesions, and suppression of SRSF1 treated by TNFα in turn suppressed the expression of IFNs^10^. Despite the potentially important role that splicing plays in the mechanism of psoriasis, analyzing alternative splicing in psoriasis has received relatively little attention in the research community. To develop a better understanding of the disease mechanism of psoriasis, this study seeks to perform an integrative analysis to reveal missing information about splicing in psoriasis that will largely complement previous gene expression analysis.

To reveal the biological functions of the alternative splicing events in psoriasis, we performed multiple sequence alignment (MSA) between the sequences of mouse and human alternative splicing events, as the conserved splicing events are more likely to play similar roles in both species^11,12^. Our analysis revealed 18 conserved exon-skipping (ES) events between mice and humans. These conserved events are potential candidates for further functional study.

To identify the candidate splicing factors (SFs) that may be key regulators of splicing disruption seen in psoriasis, we created a database—called SFMetaDB^13^—of all RNA-Seq datasets publicly available in ArrayExpress^14^ and Gene Expression Omnibus (GEO)^15^ from gain or loss function studies of SFs in mice. Using the data source from SFMetaDB, we implemented a signature comparison method to infer the critical SFs for psoriasis. The splicing changes in a psoriasis mouse model^16^ and the SF perturbation datasets were used to derive the splicing signatures. By comparing the signatures of psoriasis datasets to the splicing signatures of our splicing signature database, we revealed nine candidate SFs that potentially contribute to the regulation of alternative splicing in psoriasis. Genes regulated by such key SFs are involved in a number of critical pathways associated with psoriasis. Our large-scale analysis provides candidate targets for the biology research community to experimentally test the role of splicing in psoriasis. These results underlie the importance of completing the transcriptome landscape at the splicing level and pave the way for more detailed mechanistic studies of psoriasis in the future.

## Results

### Revealing large-scale changes in alternative splicing by analyzing RNA-Seq data from psoriasis mouse model and human skin

To investigate the role of the splicing process in psoriasis, a psoriasis mouse model was studied first to detect splicing changes. In this mouse model, the gene *Tnip1* was knocked out^16^. Notably, *TNIP1* (the homologous gene of *Tnip1)* in humans is found in a psoriasis susceptibility locus^17^. It has been shown that *Tnip1* knockout (KO) mice exhibit macroscopical psoriasis-like phenotypes, such as redness and scaling, and microscopical psoriasis-like phenotypes, such as epidermal thickening, elongated rete-like ridges, papillomatosis, retention of nuclei within corneocytes, and infiltrations with different immune cell types^16^. To reveal splicing changes, the Dirichlet Multinomial (DMN) regression^18^ was used to analyze the dataset from the *Tnip1* KO mouse model. Benjamini-Hochberg-adjusted^19^ *p-*value and the percent sliced in (PSI, Ψ) were estimated for seven types of splicing events (see Methods). Under |ΔΨ| > 0.05 and *q* < 0.05, a total of 609 differential alternative splicing (DAS) events were identified (**Table S1**). Figure 1a shows the number of DAS events for seven splicing types in the mouse model. To verify that the *Tnip1* KO mouse model recapitulated the main splicing features in human psoriasis, we performed a DAS analysis using RNA-Seq data from psoriasis patients and controls^8^. This DAS analysis identified 606 DAS events (|ΔΨ| > 0.05 and *q* < 0.05) (**Table S1**). Figure 1b shows the number of DAS events for seven splicing types in the human psoriasis dataset. In addition, Figure 2 and **Figure S1** show a few UCSC genome browser tracks of the example DAS events in *Exoc1/ EXOC1, Fbln2 /FBLN2, Fnbp1/FNBP1*, and *Atp5c1/ATP5C1* of mice/humans^20^.

**Figure 1.**
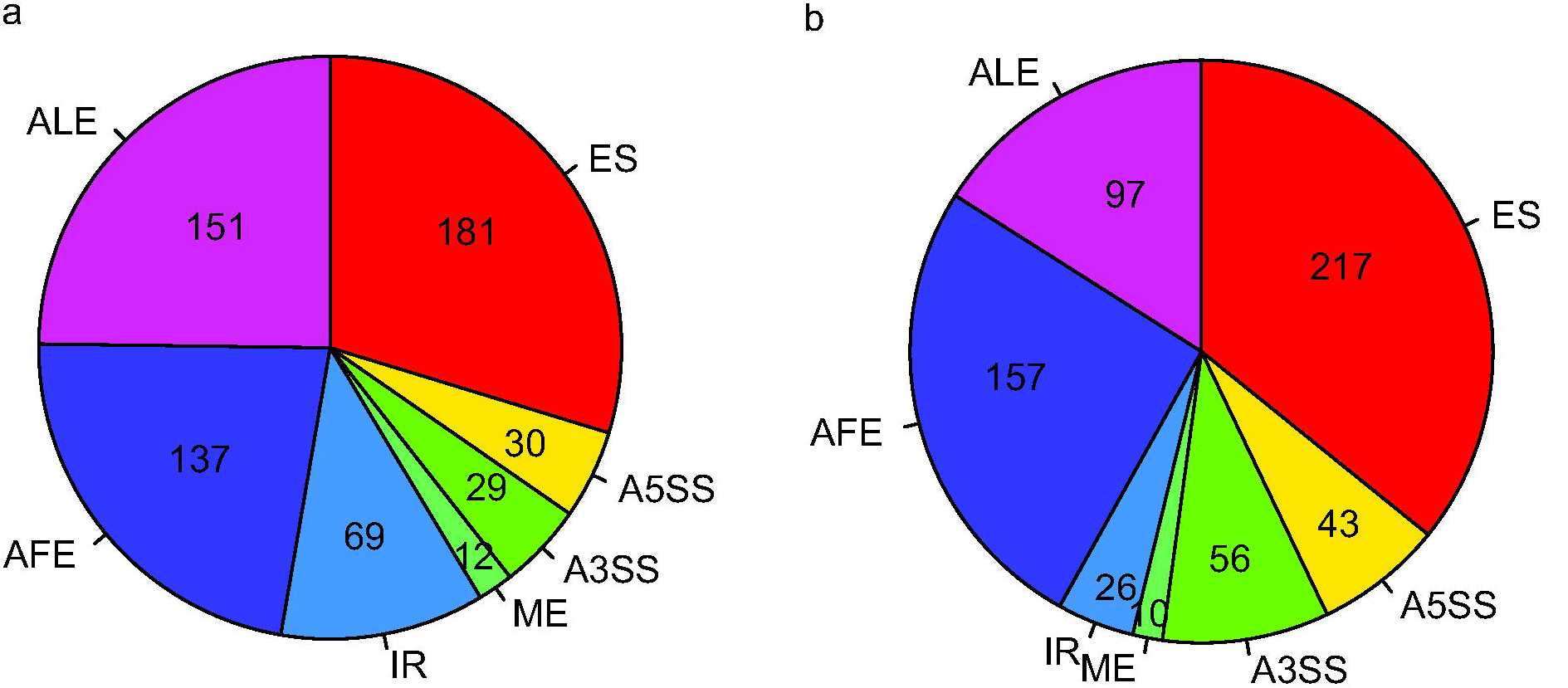
Number of DAS events for the seven splicing event types. DAS analyses were performed for the mouse and human datasets, involving seven splicing event types: ES, A5SS, A3SS, ME, IR, AFE, and ALE. Under |ΔΨ| > 0.05 and *q* < 0.05, the pie charts depict the number of DAS events for the seven splicing event types. (a) DAS analysis revealed 609 DAS events in the *Tnip1* KO mouse model dataset. (b) DAS analysis revealed 606 DAS events in the human psoriasis dataset.

**Figure 2.**
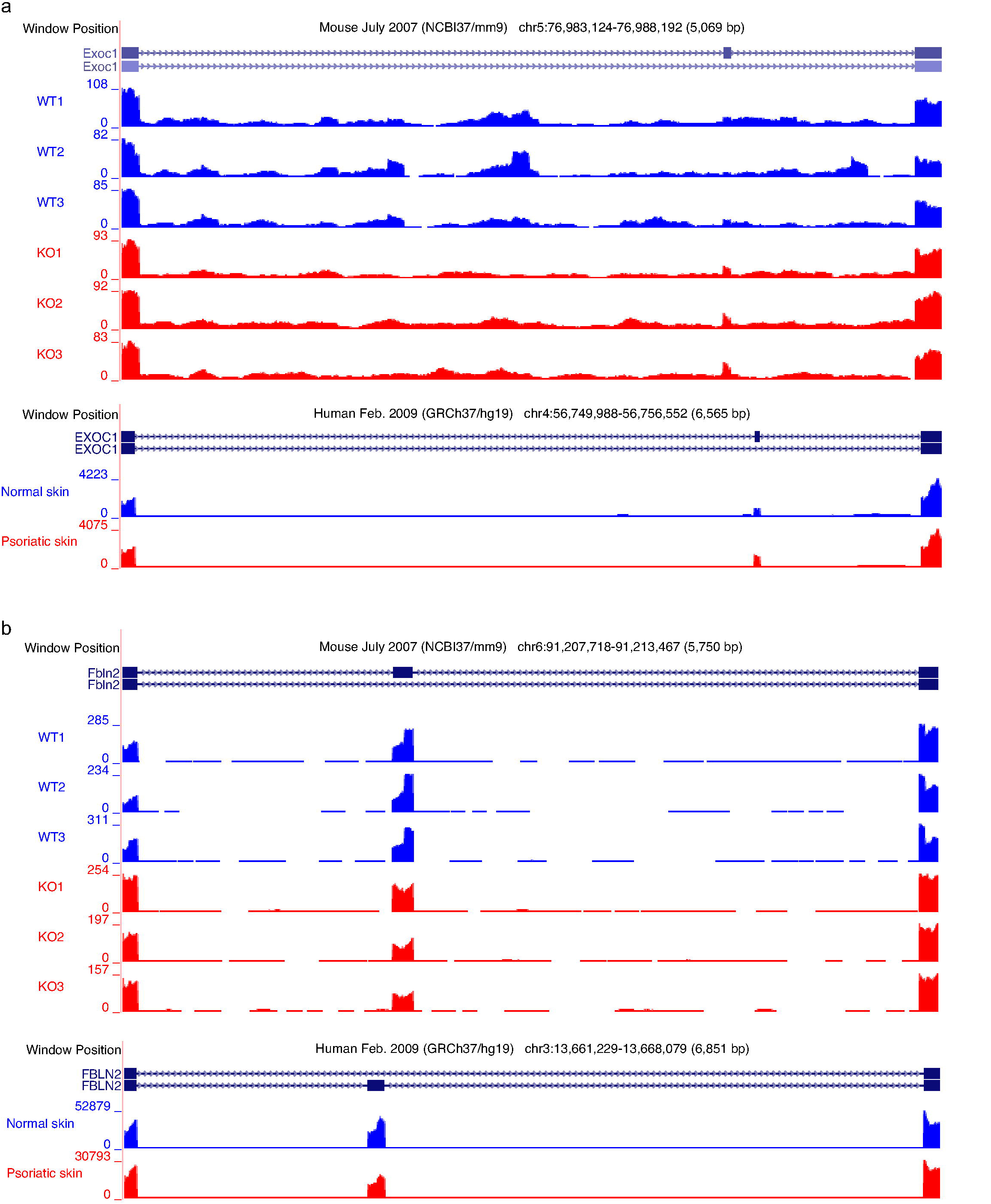
UCSC genome browser tracks visualization for the DAS events in *Exoc1/ EXOC1* and *Fbln2 /FBLN2*. The bigWigs UCSC genome browser tracks of the mapped reads in mice and humans were generated to visualize DAS events. To increase interpretability for the human psoriasis dataset, mapped reads were collapsed within groups of lesional psoriatic samples (P) and normal skin samples (N). The figures depict the expanded regions spanning over splicing events in the UCSC genome browser. (a) Visualization of the DAS event in *Exoc1/EXOC1*. The psoriatic samples have more inclusion of variable exons in both mice and humans. (b) Visualization of the DAS event in *Fbln2/FBLN2*. The psoriatic samples have less inclusion of variable exons in both mice and humans.

Our DAS results revealed many significant splicing events in the psoriasis mouse model. Figure 3 shows the heat map of PSI values for ES events in the *Tnip1* KO mouse model and in the human psoriasis dataset. In the *Tnip1* KO mouse model dataset, 64 splicing events have more inclusion of the variable exons in psoriasis, while 117 splicing events have less inclusion of the variable exons in psoriasis. In the human psoriasis dataset, 98 splicing events have more inclusion of the variable exons in psoriasis, while 119 splicing events have less inclusion of the variable exons in psoriasis. To reveal the biological functions of the genes with DAS events, gene ontology (GO) analysis (see Methods) was applied to detect the enriched GO terms for genes with DAS events in both the *Tnip1* KO mice and the human psoriasis dataset (**Figure S2** and **Table S2**). Specifically, the GO term “regulation of wound healing, spreading of epidermal cells” was enriched in both mice and humans. The wound healing process is accelerated in psoriasis, suggesting the potential role of splicing changes in psoriasis^21^. In addition, the actin-filament‒ related GO terms “negative regulation of actin filament depolymerization” and “actin filament reorganization” were enriched in mice and humans, respectively. Dysregulation of actin filament is observed in psoriatic skins, indicating that splicing changes may contribute to the formation of psoriasis^22^. Therefore, our DAS analysis discovered large-scale splicing changes in psoriasis, providing feasible and promising new features to study the role of splicing in the pathogenesis of psoriasis.

**Figure 3.**
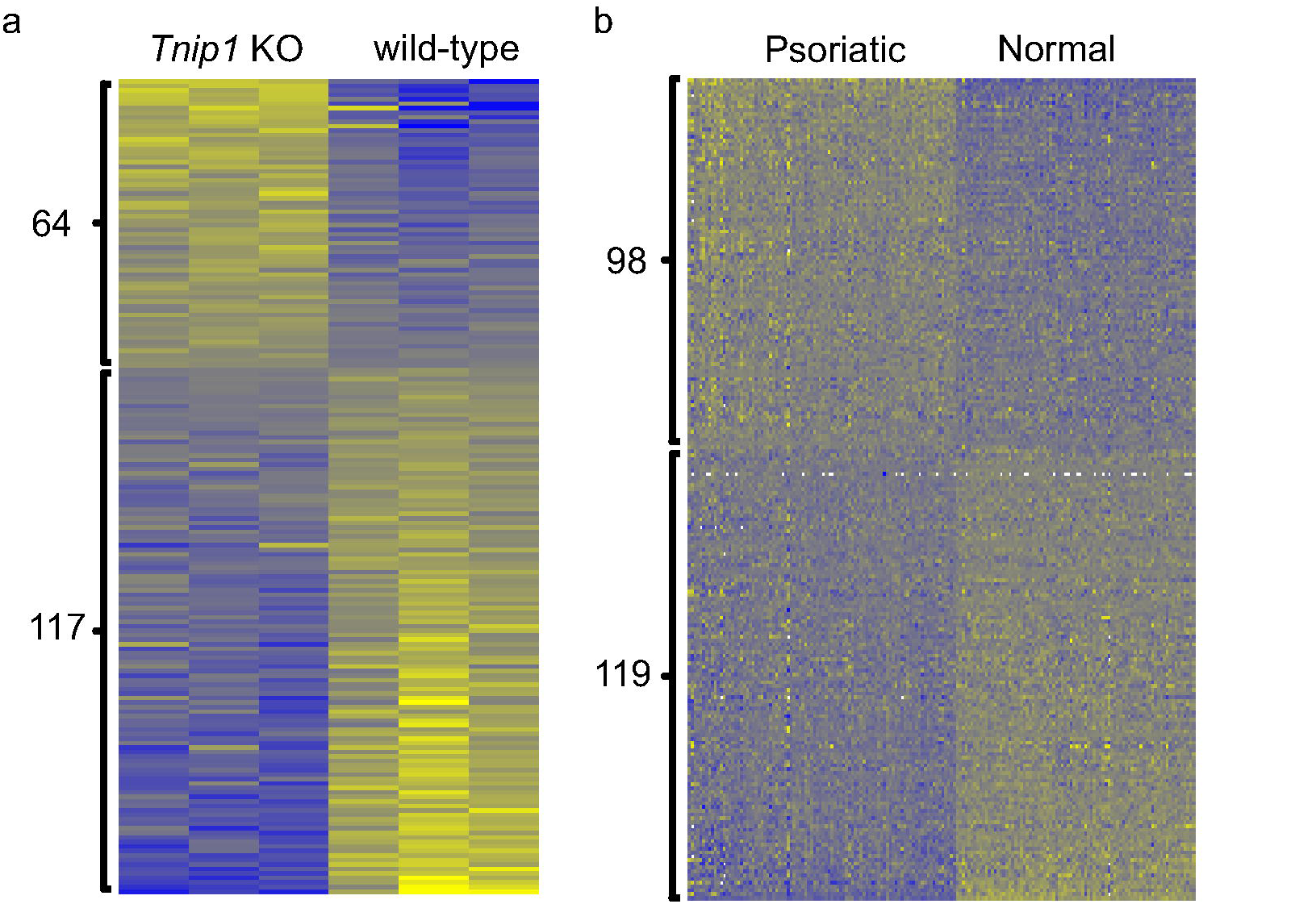
Heat map of PSI values for alternative ES events in the *Tnip1* KO mouse model dataset and the human psoriasis dataset. Yellow: high PSI. Blue: low PSI. (a) The heat map of the PSI values between three KO samples and three wild-type samples in mice. 64 of 181 ES events have more inclusion of variable exons in psoriasis, and 117 ES events have less inclusion of variable exons in psoriasis. (b) Heat map of the PSI values between 92 lesional psoriatic skins and 82 normal control skins in humans. 98 of 217 ES events have more inclusion of variable exons in psoriasis, and 119 ES events have less inclusion of variable exons in psoriasis.

### Revealing conserved splicing events in both mice and humans by splicing conservation analysis

To identify the most critical splicing changes in psoriasis, we conducted a splicing conservation analysis to reveal the splicing changes common to both the *Tnip1* KO mouse model dataset and the human psoriasis dataset. By mapping mouse and human gene symbols using HomoloGene^23^, we detected 89 homologous genes with DAS events in both mice and humans (Figure 4). The Fisher’s exact test showed significant enrichment of the common homologous genes with *p* = 1.7×10^−32^ (see Methods). This supports the conclusion that there is commonality in splicing underlying psoriasis in both mice and humans.

**Figure 4.**
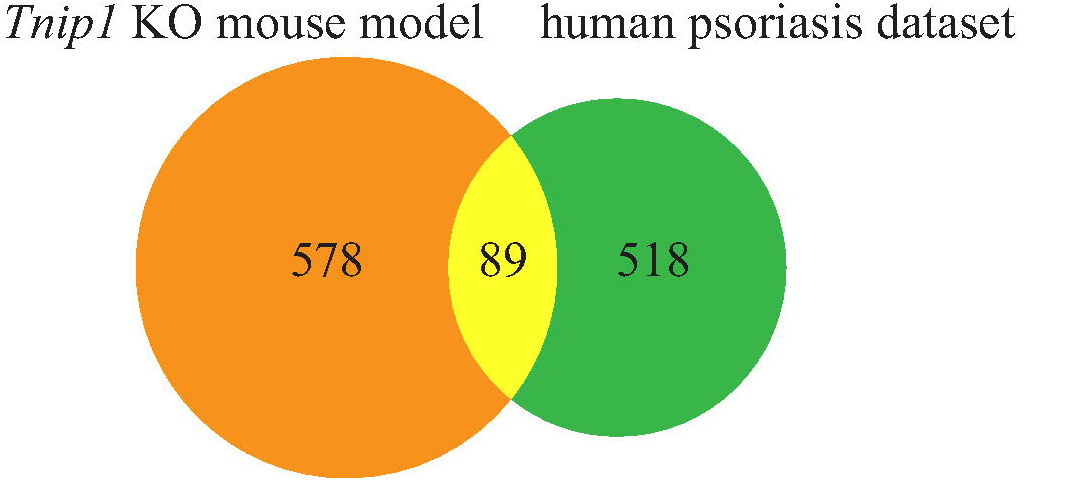
Venn diagram of the genes with DAS events in the *Tnip1* KO mouse model dataset and the human psoriasis dataset. To investigate the genes with DAS events, we ended up with 667 genes in the *Tnip1* KO mouse model dataset and 607 genes in the human psoriasis dataset. Mapping the human gene symbols to mouse homologous genes using HomoloGene resulted in 89 common homologous genes with DAS events in both species. Alternatively, 578 genes have DAS events in mice but not humans. On the other hand, 518 genes have DAS events in humans but not mice. Taking 12,233 homologous genes expressed in both species as the background genes, the Fisher’s exact test showed significant enrichment of the common homologous genes with *p* = 1.7×10^−32^.

To further characterize the conservation of splicing in mice and humans, we compared the isoform sequences between them. By the splicing conservation analysis at the isoform level (see Methods), we ended up with 24 homologous genes with conserved isoform sequences for the common splicing events in human and mouse gene annotation (**Supplemental Data S1, Table S3**). The high proportion of conserved isoform sequences for the common splicing events (24 of 33) suggested feasible and promising conservation of splicing changes between the *Tnip1* KO mouse model dataset and the human psoriasis dataset (see the representative MSA of *Exoc1/EXOC1* in **Supplemental Document S1**).

To identify the splicing features in psoriasis, we further evaluated whether the common splicing events were conserved in the same isoform between the *Tnip1* KO mouse model dataset and the human psoriasis dataset. Specifically, we checked whether the splicing events shared the same inclusion pattern of variable exons in mouse and human. We ended up with 18 alternative splicing events conserved in the same isoform, which means that the splicing events have more or less inclusion of variable exons in the same way between the two species (Table 1). The corresponding 18 homologous genes with conserved alternative splicing events include *ABI1, ARHGAP12, ATP5C1, CTTN, DNM1L, EXOC1, FBLN2, FNBP1, GOLGA2, GOLGA4, MYH11, MYL6, MYO1B, PAM, SEC31A, SLK, SPAG9*, and *ZMYND11*. The MSA results for the 18 common spliced genes can be found in **Supplemental Data S1** and **Table S3**. Of the 18 conserved splicing events, eight were largely spliced in both species, with over 10% PSI differences (Table 1). Figure 2 and **Figure 1S** show the conserved splicing events in *Exoc1/ EXOC1, Fbln2 /FBLN2, Fnbp1/FNBP1*, and *Atp5c1/ATP5C1* using the UCSC genome browser tracks^20^. Our conservation analysis identified the 18 conserved splicing events, suggesting that the splicing features in the psoriasis mouse model dataset can be recapitulated in the human psoriasis dataset, and further, the 18 conserved splicing events can be promising targets to follow to study the splicing mechanism in psoriasis.

**Table 1.**
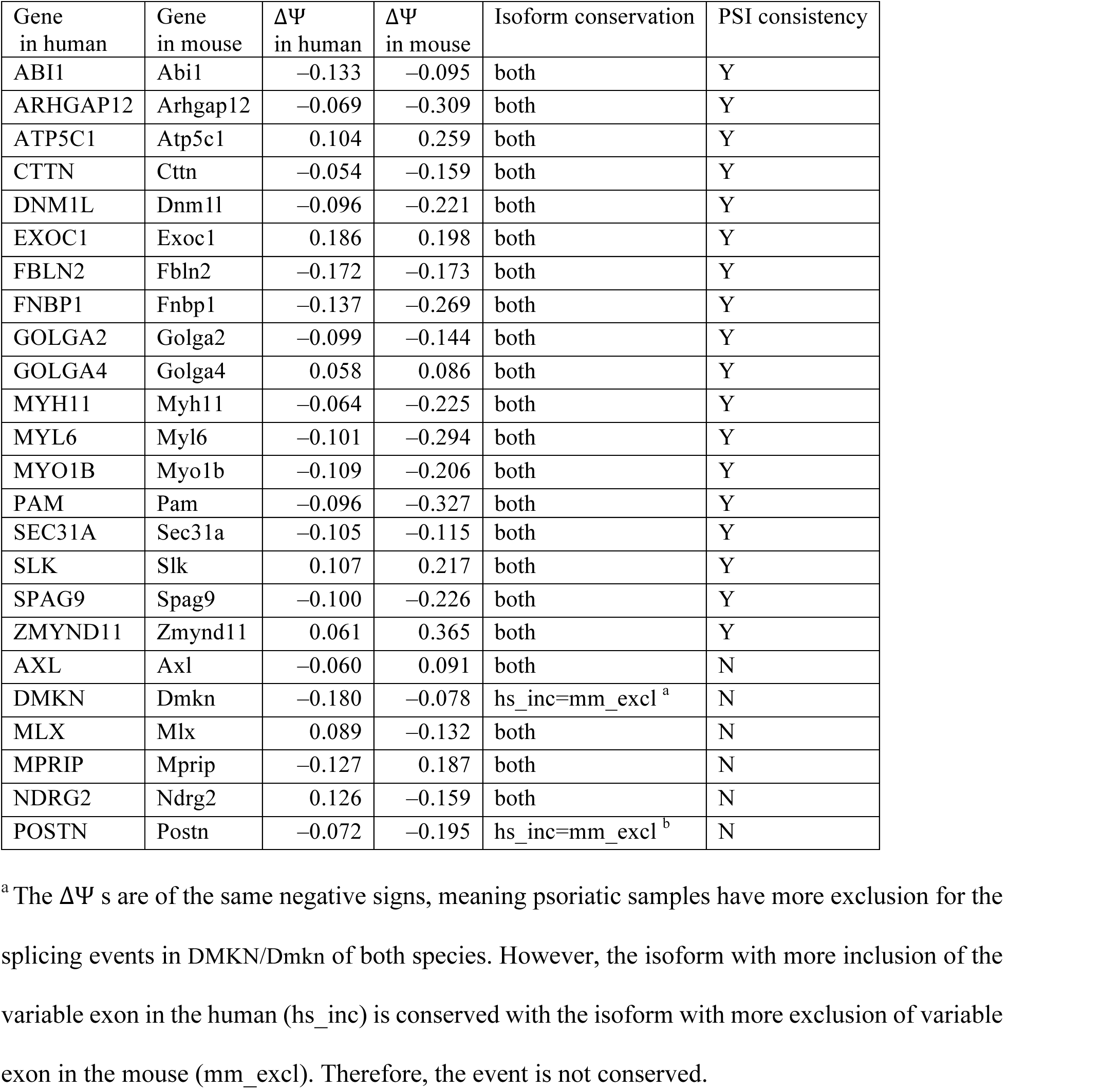
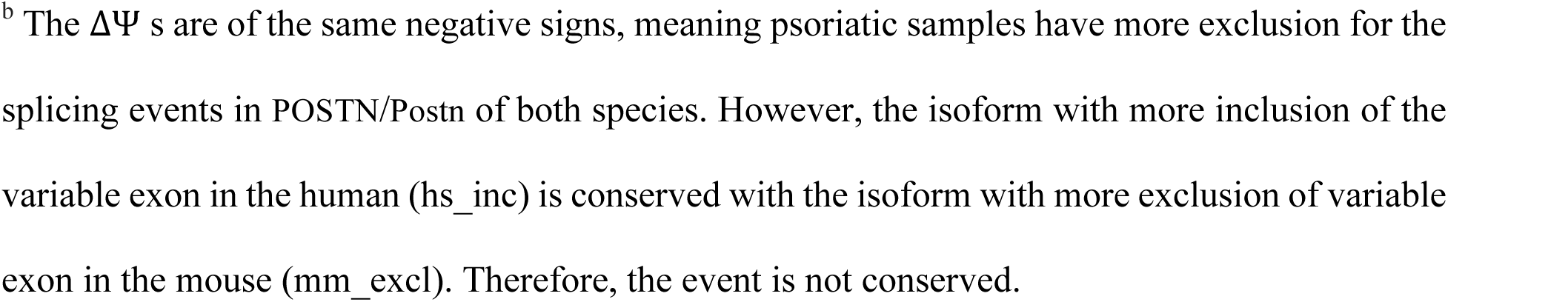
Identification of the conserved splicing events between the *Tnip1* KO mouse model dataset and the human psoriasis dataset. The column “PSI consistency” marks ‘Y’ for the 18 splicing events conserved in both mice and humans.

| Gene in human | Gene in mouse | $\Delta\Psi$ in human | $\Delta\Psi$ in mouse | Isoform conservation | PSI consistency |
| --- | --- | --- | --- | --- | --- |
| ABI1 | Abi1 | −0.133 | −0.095 | both | Y |
| ARHGAP12 | Arhgap12 | −0.069 | −0.309 | both | Y |
| ATP5C1 | Atp5c1 | 0.104 | 0.259 | both | Y |
| CTTN | Ctnn | −0.054 | −0.159 | both | Y |
| DNM1L | Dnm1l | −0.096 | −0.221 | both | Y |
| EXOC1 | Exoc1 | 0.186 | 0.198 | both | Y |
| FBLN2 | Fbln2 | −0.172 | −0.173 | both | Y |
| FNBP1 | Fnbp1 | −0.137 | −0.269 | both | Y |
| GOLGA2 | Golga2 | −0.099 | −0.144 | both | Y |
| GOLGA4 | Golga4 | 0.058 | 0.086 | both | Y |
| MYH11 | Myh11 | −0.064 | −0.225 | both | Y |
| MYL6 | Myl6 | −0.101 | −0.294 | both | Y |
| MYO1B | Myo1b | −0.109 | −0.206 | both | Y |
| PAM | Pam | −0.096 | −0.327 | both | Y |
| SEC31A | Sec31a | −0.105 | −0.115 | both | Y |
| SLK | Slk | 0.107 | 0.217 | both | Y |
| SPAG9 | Spag9 | −0.100 | −0.226 | both | Y |
| ZMYND11 | Zmynd11 | 0.061 | 0.365 | both | Y |
| AXL | Axl | −0.060 | 0.091 | both | N |
| DMKN | Dmkn | −0.180 | −0.078 | hs_inc=mm_excl<sup>a</sup> | N |
| MLX | Mlx | 0.089 | −0.132 | both | N |
| MPRIIP | Mpriip | −0.127 | 0.187 | both | N |
| NDRG2 | Ndr2 | 0.126 | −0.159 | both | N |
| POSTN | Postn | −0.072 | −0.195 | hs_inc=mm_excl<sup>b</sup> | N |

<sup>a</sup> The $\Delta\Psi$ s are of the same negative signs, meaning psoriatic samples have more exclusion for the splicing events in DMKN/Dmkn of both species. However, the isoform with more inclusion of the variable exon in the human (hs\_inc) is conserved with the isoform with more exclusion of variable exon in the mouse (mm\_excl). Therefore, the event is not conserved.
<sup>b</sup> The $\Delta\Psi$ s are of the same negative signs, meaning psoriatic samples have more exclusion for the splicing events in POSTN/Postn of both species. However, the isoform with more inclusion of the variable exon in the human (hs\_inc) is conserved with the isoform with more exclusion of variable exon in the mouse (mm\_excl). Therefore, the event is not conserved.

### Revealing candidate splicing factors regulating splicing in psoriasis by splicing signature analysis in mouse

To further elucidate the splicing mechanism in psoriasis, we conducted SF screening to discover the candidate SFs that may regulate large-scale splicing events in psoriasis. Because a great number of splicing events are discovered in mouse psoriasis datasets, we hypothesize that SFs may play critical roles in the regulation of these events. To screen for the candidate SFs, we manually curated a list of RNA-Seq datasets with gain- or loss-of-function of mouse SFs^13^ (see Methods). Using the datasets in SFMetaDB, we systematically compared the splicing changes in the psoriasis mouse model dataset with the effects of SF perturbation using a splicing signature comparison workflow (Figure 5, see Methods). Our splicing signature comparison approach screened the SF perturbation datasets related to a total 31 SFs for splicing regulators in the mouse psoriasis dataset, where nine SFs showed significant overlapping splicing changes in psoriasis, including NOVA1, PTBP1, PRMT5, RBFOX2, SRRM4, MBNL1, MBNL2, U2AF1, and DDX5 (**Table S4**), which are potential regulators responsible for splicing changes in psoriasis.

**Figure 5.**
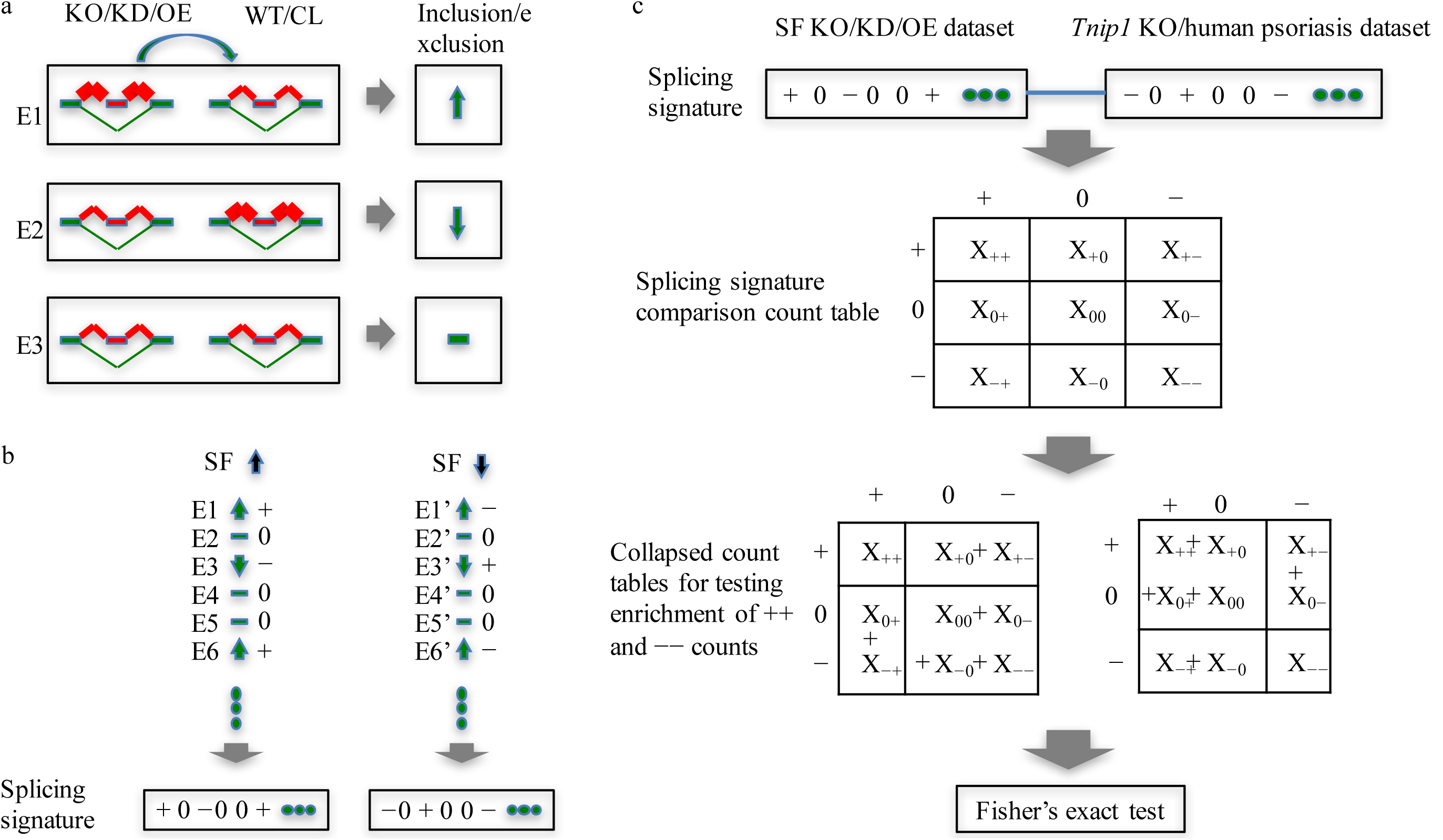
Splicing signature comparison workflow for the discovery of candidate SFs that regulate alternative splicing in psoriasis. (a) The splicing events (e.g., E1) with a more frequently included variable exon in the perturbed group than in wild-type (WT) or control (CL) are denoted by upward arrows (ΔΨ > 0.05 and *q* < 0.05). The opposite cases with ΔΨ < −0.05 and *q* < 0.05 are denoted by downward arrows (e.g., E2). The rest of the cases are considered unchanged and are denoted by horizontal bars (e.g., E3). (b) A splicing event is considered positively regulated by an SF (notated as +) when there is an increase/decrease in the inclusion of the variable exon upon the over/underexpression of the SF (e.g., E1, E6, and E3’). Alternatively, a splicing event is considered negatively regulated by an SF (notated as -) when there is a decrease/increase in the inclusion of the variable exon upon the over/underexpression of the SF (e.g. E3, E1’, and E6’). The rest of the events are considered not regulated by the SF and are notated as 0. A splicing signature is defined as the vector of +/-/0s, indicating how an SF regulates a given set of alternative splicing events. (c) Splicing signatures were first calculated for the SF perturbation (KO/KD/OE) datasets and the *Tnip1* KO mouse/human psoriasis datasets. A 3×3 contingency table was tabulated for the common splicing events between an SF perturbation dataset and the *Tnip1* KO mouse dataset or the human psoriasis dataset. This contingency table can be collapsed into 2×2 tables for testing enrichment of + + events and – - events, respectively, using Fisher’s exact test.

### Confirming the key splicing regulators in humans

To confirm the importance of these nine SFs in mice, we performed a similar splicing signature comparison analysis in humans. Using the human homologous symbols of these nine mouse SFs, we curated on GEO^15^ the human RNA-Seq datasets with these genes perturbed. Our curation resulted in four datasets for three of nine human SFs—GSE59884 ^24^ and GSE69656 ^25^ for PTBP1, GSE66553 for U2AF1, and GSE76487 ^26^ for MBNL1. The splicing signature comparison analysis (Figure 5, see Methods) using splicing signatures from the human psoriasis dataset and the four datasets of the three human SF perturbation showed significantly overlapped splicing changes between the human psoriasis dataset and the SF perturbation datasets of three human SFs— i.e., PTBP1, U2AF1, and MBNL1 (**Table S4**). These results suggest the important role of these three SFs in potentially regulating splicing in psoriasis.

### Revealing potential candidate SFs that regulate splicing in psoriasis using conserved splicing events in SF perturbation datasets

To identify the potential SFs that regulate the conserved splicing events in psoriasis, we investigated the consistency of regulation direction of splicing events in the mouse/human dataset and the SF perturbation datasets. The 18 conserved ES events were significantly conserved in the *Tnip1* KO mouse model dataset and the human psoriasis dataset, indicating the key spliced genes in psoriasis. Upon checking whether the splicing events were positively/negatively regulated by the SF in the same way in the SF perturbed datasets and the psoriasis datasets (Figure 5b), we ended up with 12 SFs (CELF1, CELF2, DDX5, MBNL1, MBNL2, NOVA1, PRMT5, PTBP1, RBFOX2, SF3A1, SRRM4, and U2AF1) potentially regulating 13 splicing events (*Abi1, Arhgap12, Atp5c1, Cttn, Exoc1, Fbln2, Golga2, Golga4, Myl6, Pam, Sec31a, Spag9*, and *Zmynd11*) in the *Tnip1* KO mouse model dataset and three SFs (PTBP1, U2AF1, and MBNL1) potentially regulating five splicing events (*ABI1, CTTN, GOLGA2, MYL6*, and *PAM*) in the human psoriasis dataset. The detailed candidate splicing regulation of SFs is shown in **Table S5**. These results show the potential SFs that may regulate splicing events in psoriasis.

## Discussion

In this work, we implemented a systems biology approach based on a large-scale computational analysis to generate a biological hypothesis about the potential role of splicing in psoriasis using psoriasis mouse and human datasets, as well as a database of RNA-Seq data with perturbed SFs. This large-scale analysis suggests 18 conserved ES splicing events in psoriasis, along with several candidate SFs, that may regulate splicing in psoriasis.

Previous studies have shown the potential roles of splicing in the pathogenesis of psoriasis. For example, an isoform of *TRAF3IP2* with a mutation may cause the formation of psoriasis. The *TRAF3IP2* gene is an essential adaptor in the IL-17 signaling pathway contributing to psoriasis. The exon-2-excluded isoform of *TRAF3IP2* loses its ability to transduce IL-17 signals to regulate downstream gene expression when there is a specific amino acid substitution at the N terminus of this isoform. The expression of this mutated isoform may promote overproduction of IL-22 and IL-17. Thus, alternative splicing with this mutation predisposes carriers to the susceptibility of psoriasis^27^.

As another example, the expression of an isoform of the *Il15* gene may inhibit the proliferation of keratinocyte in mouse skin. IL-15 is a cytokine that stimulates the proliferation of T-cells and natural killer cells. The exon-7-excluded isoform of *Il15* is minimally expressed in normal skin, but its expression can be induced by a point mutation in exon 7 of *Il15*. The expression of this isoform can reduce keratinocyte activation and inhibit infiltration of neutrophil into the dermis, producing a less-inflammatory response. Because *Il15* is within the PSORS9 psoriasis susceptibility locus, this work suggested a potential alternative splicing mechanism by which *Il15* contributes to the pathogenesis of psoriasis^28^.

Computational analysis has been shown to be a feasible and effective way to generate novel biological hypotheses. For example, a network reconstruction method was used to infer MPZL3 as a key factor in the mitochondrial regulation of epidermal differentiation^29^. As another example, a gene regulatory network analysis was applied to identify MAF and MAFB as the key transcription factors in epidermal differentiation^30^. We performed DAS analysis on the publicly available psoriasis mouse model dataset^16^. The large-scale DAS changes in the *Tnip1* KO samples compared with wild-types provided a feasible discovery of the key splicing features in psoriasis. To facilitate the discovery of the key splicing events in psoriasis, we also performed DAS analysis on a publicly available human psoriasis dataset^8^. The sharing of a large number of differential splicing events between the two species underlies the potential importance of splicing in psoriasis. The subsequent MSA analysis of the isoform sequences strengthened the discovery of the conserved splicing events in psoriasis. In addition, the splicing signature comparison workflow was applied to infer the potential candidate SFs that may regulate splicing changes in psoriasis. These computational analyses suggest the critical role that alternative splicing may play in psoriasis.

Our results suggest the potential contribution of selected splicing events in the mechanism of psoriasis. From MSA analysis, we detected 18 conserved genes sharing a common splicing pattern between the *Tnip1* KO mouse model dataset and the human psoriasis dataset, several of which can be potentially critical in psoriasis. For example, Exocyst complex component 1 (*Exoc1*) gene is a component of exocyst complex that determines the docking sites for targeting exocytic vesicles of the plasma membrane. The components of exocyst complex have been demonstrated as critical to angiogenesis^31^. As another example, Fibulin-2 (*Fbln2*) is a gene in the fibulin family that encodes an extracellular matrix protein, which plays an important role during organ development. The extracellular matrix molecules and extracellular matrix remodeling regulate angiogenesis^32^. Since angiogenesis is critical for the progression of psoriasis^33^, our results suggest a putative role of the splicing *of Exoc1* and *Fbln2* in psoriasis.

Our conservation analysis also identified that abl-interactor 1 *(Abi1)* shares a similar change in alternative splicing between mice and humans. The RNA expression of *ABI1* is universally identified in human tissues, and the protein ABI1 is expressed particularly high in human skin cells^34^. *Abi1* modulates the epidermal growth factor receptor pathway substrate 8 *(Eps8)* to regulate the remodeling of actin cytoskeleton architecture^35^. Abnormal actin cytoskeleton organization may occur in keratinocytes of lesional psoriatic skin^22^, suggesting a potential role of *Abi1* in forming psoriasis. As another identified regulator, cortactin *(CTTN)* is expressed in all human tissues, and the protein CTTN is highly expressed in human skin cells^34^. The depletion of *Cttn* represses keratinocyte growth factor receptor (KGFR) internalization and polarization, inhibiting cell migration^36^. Notably, the migration of epidermal Langerhans cells is inhibited in chronic plaque psoriasis^37^. Our analysis identified a conserved splicing pattern in *Cttn*, suggesting the potential contribution of the alternative splicing *of Cttn* to psoriasis. As another example, the splicing pattern of STE20-like kinase *(Slk)* was strongly conserved (ΔΨ > 10%) between mice and humans. Since cell migration is inhibited in psoriasis^37^, the splicing changes *of Slk* may contribute to psoriasis by aecting cell migration^38^. The above results of our conservation analysis demonstrate that splicing changes potentially affect several processes related to psoriasis. Particularly, the extracellular matrix mediates immune response^39^, the actin cytoskeleton plays a critical role in nearly all stages of immune system functions^40^, and the cell migration process belongs to innate immune cell functions^41^. Therefore, the dysregulation of these processes may result in immune dysregulation, leading to the formation of psoriasis.

In addition, our splicing signature comparison analysis identified a number of potential SF contributors to psoriasis. Despite the fact that most of them have not been previously linked to psoriasis in the literature, it has been shown that the repression of *Ptbp1* can lead to skin developmental defects^42^, highlighting that alternative splicing events regulated by PTBP1 may contribute to psoriasis. Further study is required to fully elucidate the contribution to psoriasis from the splicing-mechanism point of view.

To underline the role of splicing changes in psoriasis, GO analysis of expression changes was conducted in comparison to splicing changes. Specifically, GO analysis for up-regulated genes in mice and humans identified skin-development-related GO terms (**Figure S3** and **Table S6**). For example, the GO terms “keratinization” and “cornified envelope” were enriched for up-regulated genes in both mice and humans. The aberrant proliferation of keratinocyte contributes to the development of psoriasis^43^, and cornified envelope is involved in skin development^44^, suggesting the role of expression changes in psoriasis. However, the GO term “regulation of wound healing, spreading of epidermal cells” was uniquely identified by the splicing changes but not expression changes in both mice and humans. Due to the role of the wound healing process in psoriasis^21^, splicing changes demonstrated their particular role in psoriasis. Furthermore, an additional GO analysis was performed using alternatively spliced genes as the foreground and up-regulated genes as the background. As a result, the wound healing process was also enriched (data not shown), suggesting that the splicing changes were overrepresented in expression changes in psoriasis.

Our computational analyses suggest a potentially important role of splicing in psoriasis, which needs be validated *in vivo* or *in vitro*. A number of DAS events identified in psoriasis were linked to psoriasis supported by literature evidence, and further experimental validation of significant predictions is expected to further strengthen the support of our discovery. The 18 conserved splicing events identified in our DAS analysis can be good candidates for experimental validation. In addition, *in vivo* experiments of the identified splicing regulators may provide a better understanding of how alternative splicing is disrupted in psoriasis.

In conclusion, our DAS analysis suggests a number of splicing events related to psoriasis conserved in mice and humans, as well as SFs that may be responsible for the regulation of splicing in psoriasis. These conserved splicing events and potential candidate SFs pave the way for the research community to study the role of splicing in psoriasis. The computational DAS analysis is a feasible and efficient way to generate biological hypotheses about the role of splicing in psoriasis.

## Methods

### Differential alternative splicing analysis using RNA-Seq data

To identify the DAS events, we performed DAS analysis for the *Tnip1* KO mouse model dataset (GSE85891), where the *Tnip1* KO mice and controls were treated for two days with imiquimod (IMQ)^16^, and for the human psoriasis dataset (GSE54456), where the human lesional psoriatic and normal skins established large-scale gene expression data^8^. We first aligned the raw RNA-Seq reads to mouse (mm9) or human (hg19) genomes using STAR (version 2.5.1b)^45^ with default settings, and only uniquely mapped reads were retained for further analysis. The number of reads for each exon and exon-exon junction in each RNA-Seq file was computed by using the Python package HTSeq^46^ with the annotation of the UCSC KnownGene (mm9 or hg19) annotation^47^. DMN was used to model the counts of the reads aligned to each isoform of each event^18^, and the likelihood ratio test was used to test the significance of the changes in alternative splicing between psoriasis samples and controls^48^. We calculated the *q*-values from the *p-*values in the likelihood ratio test by the Benjamini-Hochberg procedure^19^. The DAS events are classified into seven splicing types: Exon skipping (ES), alternative 5’ splice sites (A5SSs), alternative 3’ splice sites (A3SSs), mutually exclusive (ME) exons, intron retention (IR), alternative first exons (AFEs) and alternative last exons (ALEs). In addition, PSI was used to evaluate the percentage of the inclusion of variable exons relative to the total mature mRNA in the splicing events^49^. The PSI was originally defined for ES events. Here, its definition is expanded to describe the changes in splicing of all the splicing types in our DAS analysis. Specifically, the splicing event types ES, A5SS, A3SS, ME, and IR involve two isoforms where one isoform is longer. We calculated the PSI as the percentage usage of the longer isoform compared with both isoforms. For the splicing events AFE and ALE, we calculated PSI as the percentage of usage of the proximal isoform (the isoform with the variable exon closer to the constitutive exon) relative to both isoforms of the event. The DAS events are identified under |ΔΨ| > 0.05 and *q* < 0.05.

### Gene ontology analysis

To examine the biological functions of the genes in the *Tnip1* KO mice and the human psoriasis dataset, GO analysis was performed to screen for the enriched GO terms using Fisher s exact test with the null hypothesis *H*_0_: log odds ratio < 1. In the test of enriched GO terms for the genes with DAS events, these genes were taken as the foreground, and the expressed genes were taken as the background. To reveal the enriched GO terms for differentially expressed genes, specifically up-regulated genes were taken as the foreground and expressed genes were taken as the background. The estimated log odds ratio was also retained for overlapped GO terms. Enriched GO terms were identified under *p-*value *<* 0.05.

### Splicing conservation analysis

To reveal the biological function of DAS events in psoriasis, we performed splicing conservation analysis between mice and humans. We first checked whether the homologous genes between the two species both had the DAS events. By mapping the human gene symbol to the mouse homologous gene symbol using HomoloGene^23^, we constructed a contingency table consisting of the counts of the homologous genes in both species with DAS events or in only one species with DAS events. Taking the homologous genes expressed in both mice and humans as the background genes, the Fisher’s exact test was used to test the enrichment of common homologous genes with DAS events in both species.

Additionally, we compared the isoform sequences between mice and humans. Within the 89 homologous genes with DAS events, 33 showed ES events in both species. To investigate the conservation of splicing changes in these 33 genes, we performed MSA analysis of the ES events in these genes. We first extracted the two isoform sequences that cover each of the ES events— i.e., the upstream and downstream exons in the event are included in both isoforms, but the variable exon is included in only one of the isoforms. Within each homologous gene, we compared all the mouse-human splicing event pairs. In each comparison, we constructed an MSA of the translated protein sequences or the predicted mRNA sequences of the extracted isoforms in mice and humans using MAFFT^51^. For the coding events, we constructed the MSA for the translated protein sequences. Alternatively, for the events with noncoding isoforms, we built the MSA for the predicted mRNA sequences. The MSA results between the mouse and human isoforms revealed a commonality of splicing events between mice and humans.

### Mouse splicing factor perturbation database

To screen for the candidate SFs that may regulate splicing in psoriasis, we curated a set of mouse RNA-Seq datasets with perturbed SFs. Our curated datasets were deployed as a database called SFMetaDB^13^, which hosts the full mouse RNA-Seq datasets with perturbed SFs (knocked-out/knocked-down/overexpressed). To curate the mouse SF perturbation database in SFMetaDB, we extracted 315 RNA SFs in GO (accession GO:0008380) for the mice^50^. For each SF, we used the gene symbol to search against ArrayExpress^14^ for mouse RNA-Seq datasets. For the retrieved results from ArrayExpress, we performed manual curation of the dataset to make sure the SF was perturbed in the dataset. We ended up with 34 mouse RNA-Seq datasets for the perturbation of 31 SFs. These 34 SF perturbation datasets provided the precious raw data for us to induce the candidate SFs that regulate splicing in psoriasis.

### Splicing signature‒based connectivity map

To identify the candidate SFs that regulate splicing events in psoriasis, we first determined whether the expression of SFs increased or decreased in the *Tnip1* KO mouse dataset and the human psoriasis dataset using the following procedure. The raw RNA-Seq reads were aligned to mouse/human genome using STAR, the same as the DAS analysis. The uniquely mapped reads were used to calculate the read-counts for each gene against the UCSC KnownGene annotation (mm9/hg19). A table of read-counts for all the genes and all the samples was created and normalized by DESeq^52^. The fold change calculated from this normalized count table was used to determine whether the expression of an SF increased or decreased.

Then, we checked how the splicing events were regulated by the SFs in the SF perturbation datasets by comparing these splicing events with the events in the psoriasis datasets. For example, if 1) a splicing event was positively regulated by an SF according to an SF perturbation dataset— i.e., the inclusion of the variable exon of the event was increased (Figure 5a) upon the overexpression of the SF or the inclusion of the variable exon of the event was decreased upon the knock-down/knock-out of the SF in the SF perturbation dataset (Figure 5b), and 2) the same variable exon was more included in psoriasis along with an increased expression of the SF or the same variable exon was less included in psoriasis along with a decreased expression of the SF, this consistency between 1) and 2) suggests that the event is likely regulated by the SF in psoriasis. If this consistency holds across a significantly large number of events, then the SF is likely a key factor responsible for the regulation of large-scale splicing changes in psoriasis. This consistency comparison approach was also used in CMap, a gene expression signature comparison method that has been widely used to detect the consistency between the gene expression signatures of a disease and the small-molecule or drug-treated samples^53^. Such a signature comparison method based on gene expression is powerful because some of the predictions have been validated *in vivo*^54^. However, most signature comparison approaches mainly focus on gene expression data and fail to detect fine-tuning of gene expression by splicing. To obviate the drawback in CMap, we applied asplicing signature-based comparison method using splicing changes in the SF perturbation datasets and the psoriasis datasets (Figure 5c). We first calculated the splicing signatures for the 34 SF perturbation datasets, where +/- indicates that an event is positively/negatively regulated by the given SF of the dataset and 0 indicates that no evidence exists that the event is regulated by the SF. Another signature vector made of +/-/0 was used to characterize the relation of an SF and the events in the psoriasis dataset. By comparing a signature from the SF perturbation dataset with a signature from the psoriasis data, a 3×3 contingency table was tabulated with rows and columns named +/-/0 and was used to see whether the two signatures match each other. To further check for the direction of the consistency, we collapsed the 3×3 table into two 2×2 tables so that the enrichment of + + events and – events, respectively, could be tested using Fisher’s exact test (Figure 5c). The SFs with significantly enriched + + events and – – events are the candidate SFs that regulate the splicing in psoriasis.

## Data availability

All data generated or analyzed during this study are included in this published article.

## Acknowledgments

This work was supported by startup funding to P.Y. from the ECE department and Texas A&M Engineering Experiment Station/Dwight Look College of Engineering at Texas A&M University and by funding from TEES-AgriLife Center for Bioinformatics and Genomic Systems Engineering (CBGSE) at Texas A&M University, by TEES seed grant, and by Texas A&M University-CAPES Research Grant Program.

## Author Contributions

J.L. carried out the major analyses. P.Y. supervised the analyses. J.L. and P.Y. wrote the manuscript. Both authors reviewed and approved the final manuscript.

